# Optimal foraging for simple organisms, the single-input marginal value theorem

**DOI:** 10.1101/2025.04.04.647000

**Authors:** Tom Orjollet–Lacomme, Roger Lloret-Cabot, Jonathan Friedman, Alfonso Pérez-Escudero

## Abstract

Most organisms devote a great deal of time and energy to locating and consuming food, making efficient foraging crucial for their evolutionary success. Consequently, Optimal Foraging has been widely studied, but mainly in relatively complex animals with strong sensory and cognitive abilities. However, most living organisms, from bacteria to lower invertebrates, lack these abilities and still must forage efficiently. Here we extend one of the main theorems in Optimal Foraging, the Marginal Value Theorem, to organisms with minimal sensory and cognitive abilities; in particular, we consider individuals whose only sensory input is their feeding rate. To do so, we reformulate a continuous version of the Marginal Value Theorem, finding that individuals should adapt their speed to achieve a marginal feeding rate equal to the average feeding rate for the environment whenever possible, where the marginal feeding rate is defined as the feeding rate they would have if they returned to an already-visited location. We also find that, when faced with uncertainty, moving intermittently is more efficient than moving at constant speed, because this speed variability enables more robust estimates of the marginal feeding rate. Our results generalize Optimal Foraging Theory to organisms with strong sensory constraints, and unify classical results derived for patchy environments to those applicable in arbitrary environments.

---

Obtaining food is not only key for survival; it is one of the most time-consuming activities across the tree of life. For this reason, even minute efficiency gains can provide a significant advantage and foraging must be highly optimized. This high degree of optimization, together with the fact that its outcome is relatively easy to measure (in terms of the amount of food collected per unit time) makes foraging an ideal test-bed to understand how organisms make decisions and execute them. For this reason, foraging is deeply studied, both from a theoretical and from an experimental point of view [40]. In particular, Optimal Foraging Theory studies the optimal strategies that animals should follow to maximize their food intake [12, 24, 35, 40].

One of the key results in Optimal Foraging Theory is the Marginal Value Theorem, which predicts the amount of time an animal must exploit a discrete food source (or food patch) before moving on to search for a fresh one [13]. Assuming that each food patch gives diminishing returns (i.e. the animal’s feeding rate decreases over time due to depletion of the food in the patch), this theorem predicts that animals should leave a food patch even if there is still some food left in it. The exact moment when they should leave is given by the time in which the instantaneous feeding rate at the food patch reaches the average feeding rate in the environment [13].

This theorem has interesting implications: When feeding on a food patch, individuals must consider not only its quality, but also the overall richness of the environment. This assessment of the environment often has a high exploration cost and requires relatively advanced cognition, so implementation of the Marginal Value Theorem reveals a sophisticated level of behavioral optimization. The predictions of the Marginal Value Theorem have been tested in many species, finding a range of results, from excellent quantitative agreement to no agreement (reviewed by [31]). It has also been tested in contexts other than foraging but sharing the same mathematics, such as mating duration in dungflies [33] and plant root growth [29]. From a theoretical perspective, the Marginal Value Theorem has been extended to situations with greater uncertainty [27, 32], continuous environments [2–4], situations in which the statistics of the environment must be learned over time [28], and more [17, 34].

Despite this intense study, the Marginal Value Theorem has been applied mostly to relatively complex organisms, mainly animals from insects upwards [31]. These organisms have advanced sensors, and in particular vision, so they can quickly scan large portions of the environment and judge its richness (for example, they can estimate instantaneously the number of food patches visible from their current position and their distance). They also have relatively large nervous systems, capable of storing and processing large amounts of information. In contrast, most living organisms, from motile bacteria to lower invertebrates, must forage without these advanced sensory and cognitive abilities. The exploration and foraging strategies of these organisms are well studied outside the Marginal Value Theorem, both experimentally and theoretically. The main framework to study their movement is random walks [9, 14, 41, 42], which are modulated in various ways in response to instantaneous sensory inputs and past experiences [6– 8, 10, 11, 16, 18, 20, 22, 36, 38]. However, to date, no version of the Marginal Value Theorem has been developed to accommodate the constraints of simple organisms.

Here we adapt the Marginal Value Theorem to organisms with minimal sensory and cognitive abilities. In particular, we consider the extreme case of an individual whose only sensory input is its feeding rate. Given that an individual with such strong sensory constraints cannot know whether the environment is patchy, we start from a continuous version of the Marginal Value Theorem that can be applied to any environment [4]. This paper is organized as follows: We first reformulate this model introducing the concept of *marginal feeding rate*, which unifies its results with those of the classical (patch-based) Marginal Value Theorem [13]. Then, we show how it can be implemented with a single sensory input (the feeding rate), but with a fragile method that requires significant computational power. Then, we show how robust near-optimal behavior can be achieved with a single sensory input and minimal computational power. Finally, we show that our results are compatible with an organism that learns the statistics of the environment over time.

## The model

Our model aims to describe an individual that only has access to local information: It can estimate the food density at its current location, but it has no information about any other point in the environment. Therefore, in contrast with visual animals that can instantaneously build a map of their surroundings, our individual has no overview of the distribution of food. Given these strong sensory limitations, an individual moving in 2D or 3D (Figure 1A) experiences the environment as a sequence of food densities (Figure 1B). Using only this information, the individual must choose its speed and direction at every point. However, here we consider only changes in speed and ask: Given a certain trajectory followed by the individual, what is the optimal speed at every point in order to maximize the overall feeding rate?

**Figure 1:**
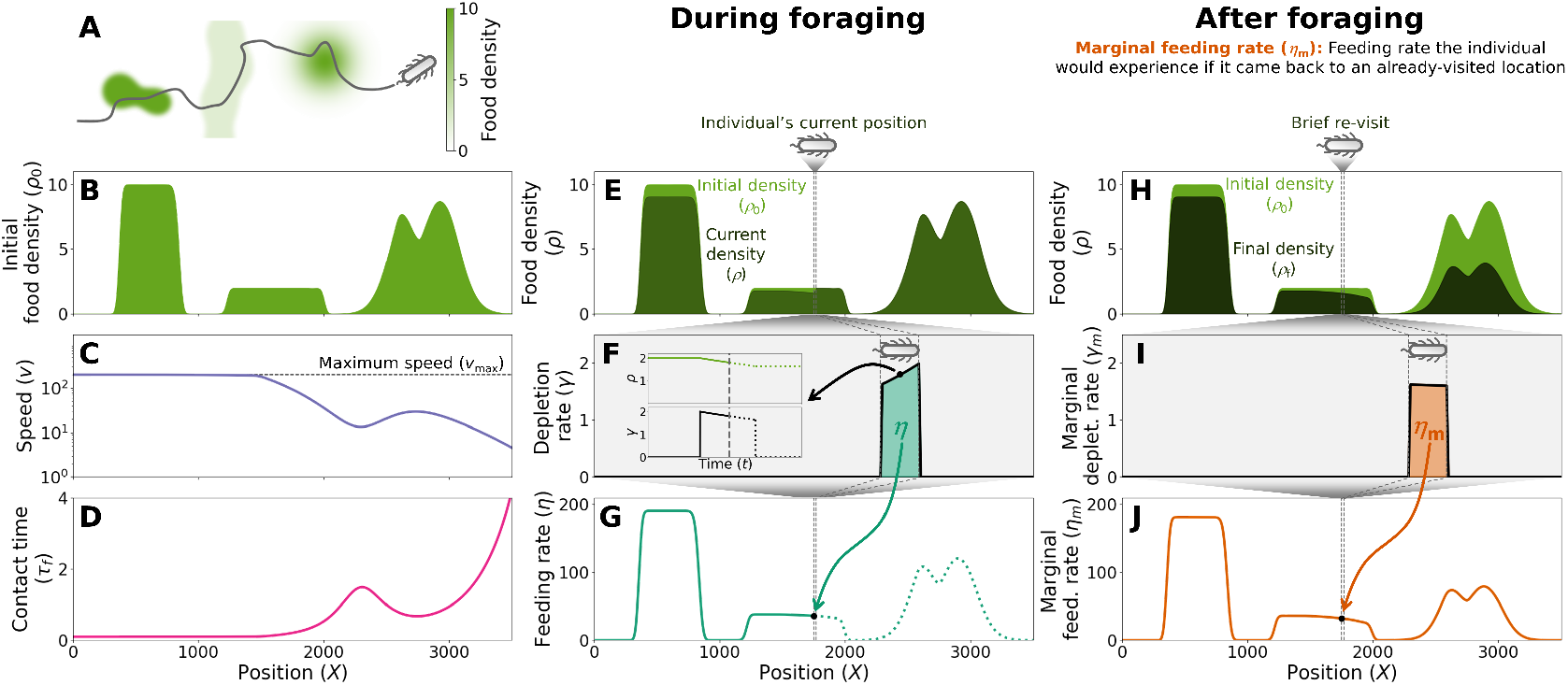
Elements of the model. **A**. Trajectory of an individual in a 2D environment. **B**. Food density encountered by the individual at each point along its trajectory. We call this density “initial density” because it is the density before depletion. **C**. Speed at each point along the individual’s trajectory. Dashed line: Maximum speed (*v*_max_). **D**. Time that each point spent in contact with the individual, once it has traversed the whole trajectory. **E**. Initial food density (light green) and food density at the current time (dark green). Vertical dashed lines: Region occupied by the individual at the current time. **F**. Depletion rate (*γ*) at each point, for a small region around the individual depicted in E. The area under the curve is the individual’s current feeding rate (*η*). Inset: Depletion rate (*γ*) and density (*ρ*) as a function of time for the point indicated by the dot in the main panel. Dashed vertical line: Current time. **G**. Feeding rate (*η*) experienced by the individual at each point. Vertical dashed lines: Region occupied by the individual at the current time. Dot: Feeding rate at the current time (equal to the area under the curve in F). **H**. Initial food density (light green) and final food density (dark green) at each point along the trajectory. Vertical dashed lines: Region occupied by a hypothetical individual returning to an already-visited location. **I**. Same as F, but for the hypothetical individual. In this case the depletion rate is computed from the final density, and we call it marginal depletion rate (*γ*_m_). **J**. Marginal feeding rate (*η*_m_) at each point along the trajectory.

To answer this question, we used a one-dimensional model, in which the animal encounters a certain food density at every point along its trajectory (Figure 1B) and must choose a strategy, defined by its speed at every point, *v*(*X*) (Figure 1C). We assume that the speed is always positive (i.e. the individual never comes back). We also assume that *v*(*X*) cannot be strictly zero, because this could create an ambiguity in its definition (the same point would have speed 0 while the individual is stopped, and a positive speed when it resumes its movement). But *v* can be arbitrarily small, so there is no practical difference with having *v* = 0. Also, this speed has an upper limit, *v*_max_, given by the individual’s physical constraints.

It’s important to note that, in contrast with the classical Marginal Value Theorem [13], our model does not assume a patchy environment: The food can be distributed in an arbitrary way, with discrete patches, smooth changes in food density, or a combination of both (Figure 1B). This generality is a key requirement to describe simple organisms, because their sensory constraints prevent them from knowing whether the food distribution is patchy, and therefore from making decisions such as “leave a food patch”. For this reason, our model follows previous works to apply the Marginal Value Theorem to non-patchy environments [4].

We assume that the individual consumes food from every point within a finite region of size *r*. The actual meaning of this region depends on the species: For an animal, it may represent the mouth; for a bacterium it may represent the whole bacterium, plus the region of space from which nutrients diffuse almost instantaneously to it; for a group of animals moving together, it may represent the whole extent of the group. While the actual meaning of this region depends on each organism, for simplicity we will refer to it as the region in contact with the individual, and to *r* as the individual’s length. We define the position of the individual at its back end, so when the individual is at *X*, it feeds from every point in the region between *X* and *X* + *r*. Once we know the individual’s length *r* and the strategy *v*(*X*), we can compute the total amount of time that each point along the trajectory spends in contact with the individual, which we call *τ*_f_(*X*) (Figure 1D).

At every point in contact with the individual, food density decreases at a rate *γ*, which we call the depletion rate (Figure 1E,F). We will illustrate our results with the simple assumption that the individual consumes a fraction of the available food in every time step, so depletion rate is simply proportional to food density, *γ* = *kρ* (Figure 1E,F). Because of this relationship between food density and depletion rate, food density decreases exponentially during the time a point is in contact with the individual (Figure 1F, inset). However, in general, the depletion rate may depend on food density in more complex ways, the only requirement being that depletion rate must decrease as food gets depleted, providing diminishing returns. In general, depletion rate may also depend on the individual’s speed, because most organisms feed less efficiently when moving fast than when moving slow. While our results are applicable for a speed-dependent depletion rate (see proof in Methods), here we will illustrate them without such dependence.

The optimal strategy is the one that maximizes the individual’s average feeding rate 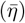, defined as the total amount of food consumed by the individual (*R*) over the whole time available for foraging (*T*), 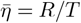.

## The optimal strategy

### The optimal strategy levels the marginal feeding rate at every point

Our key insight is the definition of the marginal feeding rate (*η*_m_), which is the feeding rate that the individual would experience if it came back to an already-visited point for an infinitesimal amount of time, and with the same speed it had when it visited the point the first time. To understand the marginal feeding rate, we first must define the actual feeding rate experienced by the individual during foraging (*η*). This feeding rate is equal to the integral of the depletion rate (*γ*) over the individual’s whole length (Figure 1F,G). The marginal feeding rate (*η*_m_) is computed in the same way, but using the depletion rate corresponding to the final food density (Figure 1H-J).

We have found that the optimal behavior is to achieve the same marginal feeding rate 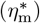 in every point wherever possible, and to move at the maximal speed at points where the initial food density is too low to achieve the target marginal feeding rate (Figure 2 A-C).

**Figure 2:**
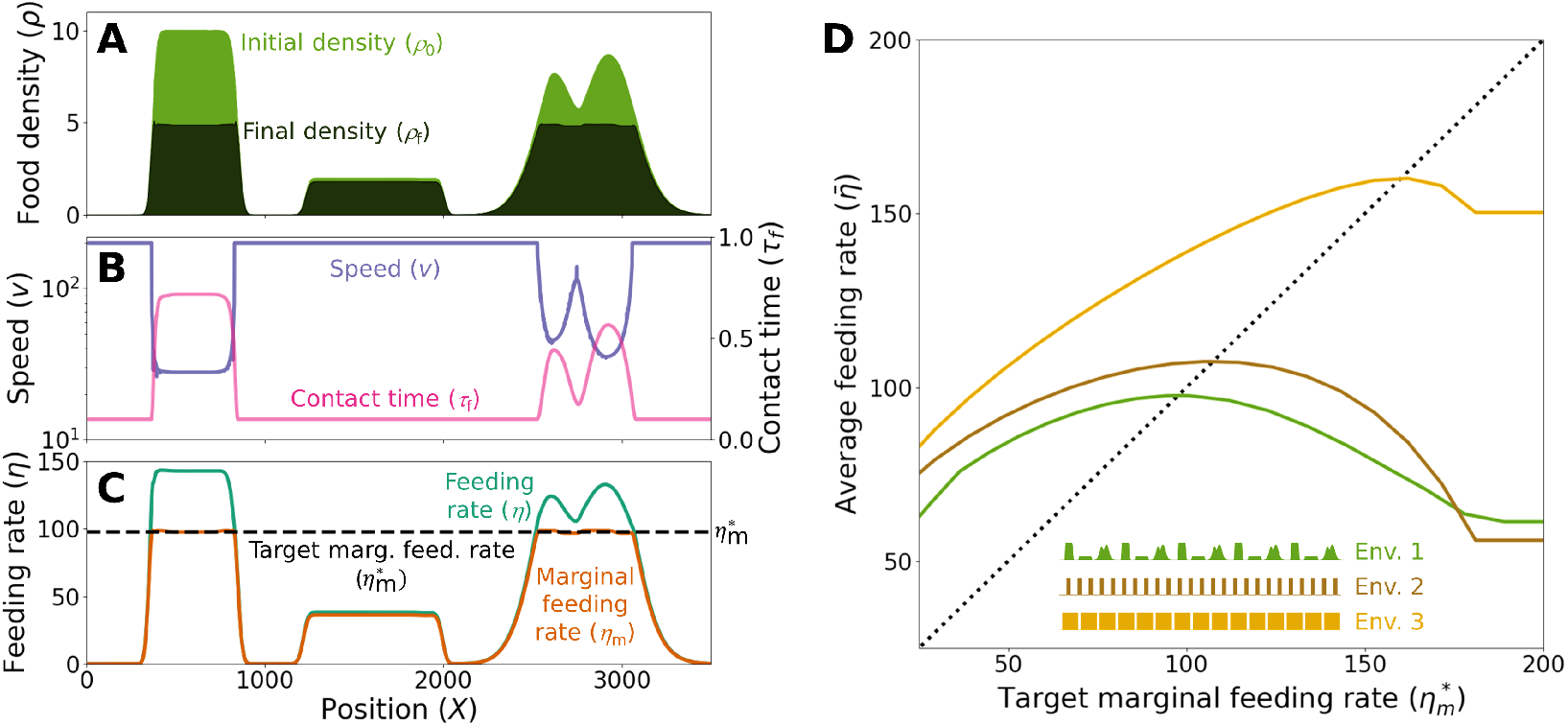
The optimal strategy levels the marginal feeding rate. **A**. Initial food density (*ρ*_0_, light green) and final food density (*ρ*_f_, dark green) at each point. **B**. Optimal speed at each point (blue), and the corresponding time that each point spends in contact with the individual (pink). **C**. Feeding rate experienced at each point, *η* (green), and marginal feeding rate, *η*_m_ (orange), for the optimal behavior. Dashed line: Target marginal feeding rate 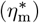 **D**. Long-term average feeding rate 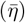 in three different repetitive environments, as a function of the target marginal feeding rate 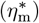. Note that the maximum of all lines fall on the 1:1 line (black dashed line). Inset: A portion of the environment represented by each color.

The intuition behind this result is relatively simple: The marginal feeding rate at point *X* turns out to be proportional to the additional amount of food that the individual could extract from that region by reducing its speed infinitesimally to stay near *X* for a bit longer (see Lemma 1 in Methods). Likewise, it is equal to the amount of food that would be lost by speeding up infinitesimally to spend a bit less time near *X*. Therefore, if after executing a given strategy, some point *X*_*A*_ has a marginal feeding rate higher than another point *X*_*B*_, then the individual would have benefited from spending more time at *X*_*A*_ at the expense of spending less time at *X*_*B*_. This transfer of time from *X*_*B*_ to *X*_*A*_ would give us an alternative strategy that extracts more food than the original one in the same amount of time, so the original strategy could not be optimal. Therefore, for the optimal strategy any two points *X*_*A*_ and *X*_*B*_ must have the same marginal feeding rate. The only exception to this argument is the case in which the individual is moving at its maximum speed at *X*_*B*_, because in this case it cannot spend less time there. Therefore, for the optimal strategy, all points where *v < v*_max_ must have the same marginal feeding rate, which we call target marginal feeding rate 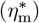, and all points where *v* = *v*_max_ must have a marginal feeding rate lower or equal to 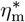 (see Methods for a formal proof).

This result is similar to previous models for continuous environments, which concluded that the individual must reach the same final food density at every point unless it’s moving at maximum speed [2–4]. Indeed, in most cases, points with the same final food density also have the same marginal feeding rate, so both results are equivalent (Figure 2 A,C). However, our result is more general and makes it easier for a forager to learn the optimal threshold, as we will show in the following section.

### The optimal target marginal feeding rate equals the average feeding rate

In the previous section we saw that the individual must target the same marginal feeding rate 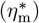 in every point, except for those in which it is moving at maximum speed. We now ask what is the optimal value of this target marginal feeding rate 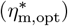. To answer this question we need to add one assumption, which was already present in the classical Marginal Value Theorem [13], called “repetitive environment”: We assume that the environment can be subdivided into many similar segments, so that the average feeding rate 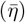 is approximately equal in each segment.

In such a repetitive environment, the optimal target marginal feeding rate is equal to the long-term average feeding rate,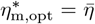. The intuition behind this result is as follows (see Methods for a formal proof): Consider the optimal strategy, whose target marginal feeding rate is 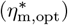. The individual can choose to increase 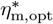 slightly, leading to an overall increase of speed, which would save an amount of time Δ*t* across the whole trajectory. This time can then be used to exploit a new segment of the environment, and in a repetitive environment this would provide an amount of food equal to 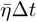. In exchange, the increase in speed would cost an amount of food equal to 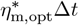 (see Lemma 1 in Methods). If the original strategy was optimal, the new strategy cannot collect more food, so we must have 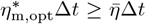. We can make the same argument for an overall decrease of speed, so for the optimal strategy we must have 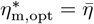.

We confirmed this result via simulations in several repetitive environments, finding that indeed the optimal value of 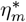 always matches the average feeding rate (Figure 2D).

### Parallel with the classical Marginal Value Theorem

Our model parallels the classical (patch-based) Marginal Value Theorem, which shows that the optimal time to leave a food patch is when the instantaneous feeding rate reaches the long-term average of the environment [13]. In the classical model, the feeding rate an animal experiences just before leaving a food patch is the same as it would experience if it came back to it, so it is equivalent to our marginal feeding rate. Therefore, the classical rule “leave the food patch when the (marginal) feeding rate matches the average feeding rate of the environment” becomes “adjust your speed so that the marginal feeding rate matches the average feeding rate of the environment” (Figure 3).

**Figure 3:**
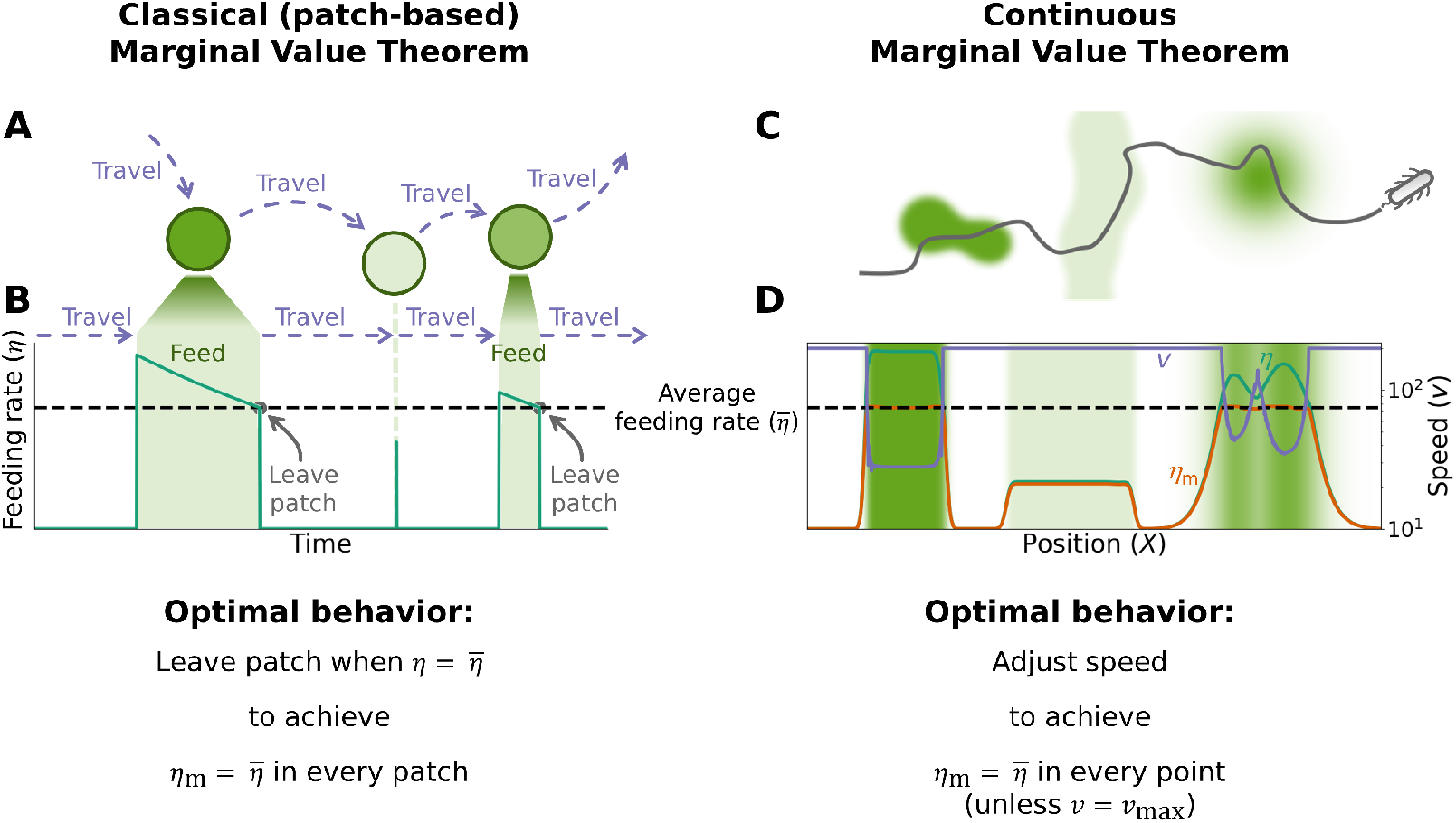
The continuous Marginal Value Theorem parallels the classical (patch-based) version. **A**. Schematic of an individual in a patchy environment. **B**. Feeding rate over time for an individual implementing the Marginal Value Theorem in a patchy environment. Dashed line: Long-term average feeding rate. Green shaded regions represent the time when the individual is feeding at a patch. The individual leaves a patch when its instantaneous feeding rate reaches the long-term average. The second patch is skipped, because its feeding rate is below the threshold from the beginning. **C**. Schematic of an individual moving in a continuous environment. Green intensity represents food density (see colorbar in Figure 1A). **D**. Feeding rate (green line), marginal feeding rate (orange line) and speed (blue) for an individual implementing the optimal behavior in a continuous environment. Background shade of green represents the initial density at each point (see colorbar in Figure 1A). Dashed line: Long-term average feeding rate in this environment.

## Near-optimal strategies with minimal sensory and cognitive abilities

The previous sections have identified the optimal behavior, but have not described how to compute it. This computation is often hard, and requires the individual to know the food density at locations outside its length. This is unrealistic, especially for simple organisms, so we will now discuss how an individual can estimate the optimal speed at every point with limited sensory inputs and computational abilities. In particular, we consider an individual that can sense the environment through a single sensory input: its instantaneous feeding rate, *η*. This feeding rate provides an estimate of the food density, but this estimate is blurred for two reasons: First, feeding rate changes non-linearly with density. Second, feeding rate is integrated across the whole individual’s length, so the individual is blind to any environmental variations at a smaller scale, and has no information about the food density at any point outside its current position.

Another challenge to find the optimal behavior is that in most cases the individual never experiences a feeding rate that equals the marginal feeding rate: Its actual feeding rate always comes from a region that is incompletely depleted (Figure 4A), and to actually experience the marginal feeding rate it would need to come back to a fully-depleted one.

**Figure 4:**
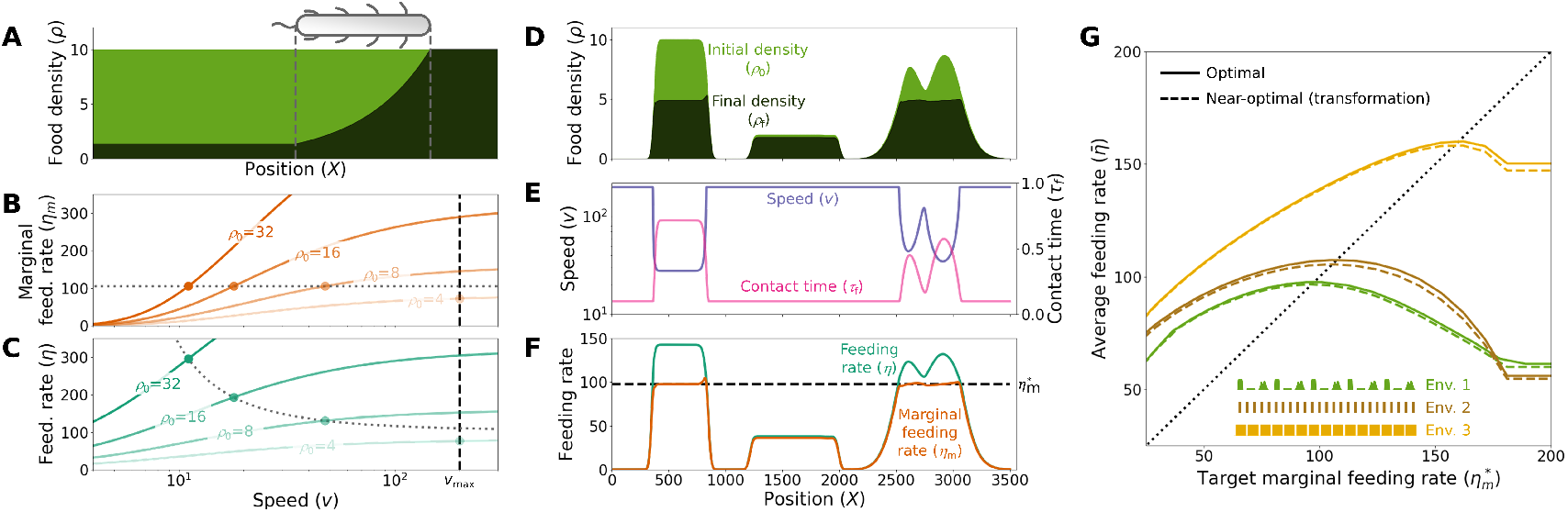
Near-optimal behavior can be achieved with a single sensory input. **A**. Initial food density (*ρ*_0_, light green) and final food density (*ρ*_f_, dark green) at each point, for an individual moving at constant speed in a uniform environment whose initial food density is *ρ*_0_ = 10 in every point. The vertical dashed lines indicate the region of space currently occupied by the individual. **B**. Marginal feeding rate (*η*_m_) as a function of speed, for the situation depicted in A, and for different values of the initial food density, *ρ*_0_ (lines follow Equation 30). Horizontal dotted line: Target marginal feeding rate 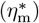. Vertical dashed line: Maximum speed (*v*_max_). Dots: Optimal speed for each initial density. **C**. Same as B, but for the feeding rate (Equation 32). Dotted line: Target feeding rate (Equation 36). **D**. Initial food density (light green) and final food density (dark green) at each point along a trajectory in a heterogeneous environment, for an individual performing near-optimal behavior. **E**. Speed (blue) and time in contact (purple) for the near-optimal behavior in the environment depicted in D. **F**. Feeding rate (blue) and marginal feeding rate (purple) achieved in every point for the near-optimal behavior in the environment depicted in D. **G**. Long-term average feeding rate 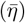 in three different repetitive environments, as a function of the target marginal feeding rate 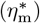. Solid lines: Optimal behavior with no sensory constrains. Dashed lines: Near-optimal behavior. Inset: A portion of the environment represented by each color.

We will first see how it is theoretically possible to approximate the target marginal feeding rate with these sensory constraints. However, this approximation requires knowing how the depletion rate depends on food density and speed, *γ*(*ρ, v*). In real conditions, this dependency may change from one food source to another, and the individual may not have enough information to infer it accurately. In this case, the feeding rate cannot be used to estimate the food density nor to find the optimal speed. However, we will see how the individual can still easily implement a near-optimal behavior by modulating its speed to get direct samples of the marginal feeding rate.

### Near-optimal behavior with a single sensory input

To find a control rule that achieves near-optimal behavior with limited sensory information, we will consider a case in which the initial food density, *ρ*_0_(*X*), and the speed *v*(*X*) are roughly constant along the individual’s length (*r*). In such a situation, the final food density left behind the individual will also be equal to a constant, *ρ*_f_ (Figure 4A), which determines the value of the marginal feeding rate. In turn, this final density (and hence the marginal feeding rate) is a function of the initial density (*ρ*_0_) and the speed (*v*): For a given value of the initial density, higher speeds lead to less depletion and hence a higher value of the marginal feeding rate (Figure 4B). For any given initial density, the optimal speed is the one that makes the marginal feeding rate *η*_*m*_(*X*) equal to the target marginal feeding rate 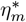 (Figure 4B, dots). If the initial density *ρ*_0_ is too low, the marginal feeding rate can never reach the target and the individual must move at its maximum speed.

One difficulty in implementing this rule is that the individual cannot measure the marginal feeding rate. It can however compute it from the actual feeding rate and the speed, which are both known to the individual (see Methods). We can thus transform the target marginal feeding rate 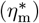 into a target feeding rate, which now depends on speed (*η*\*(*v*)) (dotted line in Figure 4C). This speed dependence is one of the main differences between the classical patch-based Marginal Value Theorem [13] and its continuous versions (such as our model and the one in [4]): In the classical patch-based Marginal Value Theorem, the threshold in feeding rate to leave a food patch is the same for every food patch, regardless of its initial density. In contrast, in the continuous versions of the Marginal Value Theorem, the target feeding rate (*η*\*) depends on the initial density (Figure 4C); only the target marginal feeding rate 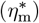 is the same for every point in space, regardless of its initial density (Figure 4B).

Once we have computed the target feeding rate for each speed (*η*\*(*v*)), the individual must simply measure its feeding rate at each time and adjust its speed to make it equal to the target. This control rule leads to the optimal speed, using the current speed (*v*) and the current feeding rate (*η*) as its only inputs. This estimate is exact only in a constant environment and moving at a constant speed, so the speed may deviate significantly from the optimal value when food density changes abruptly, such as at the edge of sharp food patches (Figure 4D-F). But its overall performance is generally good, providing a good approximation to the actual optimal behavior across a wide range of environments (Figure 4G).

However, this rule has two important limitations: First, the transformation between marginal feeding rate and feeding rate is relatively complex and leads to a speed-dependent target feeding rate, which may be hard to implement for a simple organism. Second, making the transformation from marginal feeding rate to feeding rate requires knowing precisely how the depletion rate depends on density and speed (i.e. we need to know the shape of the function *γ*(*ρ, v*)). This function is not universal, and even the same organism may experience different depletion functions in different contexts, for example due to differences in the type and state of the food. Incorrect assumptions regarding *γ*(*ρ, v*) may result in a speed that is very far from the optimal (Figure S1). The next section will present an alternative method that solves these limitations.

### Intermittent movement achieves robust near-optimal behavior

The key difficulty in implementing the optimal behavior is that the individual never experiences the marginal feeding rate (*η*_m_), as it never feeds from a region that has reached its final food density. Instead, a continuously moving individual feeds from an unequally depleted region, whose density ranges from the initial density at the front to the final density at the back (Figure 4A).

This issue can be alleviated by moving intermittently, rather than at constant speed: Consider an individual that moves at sufficiently high speed so that food depletion is negligible during movement, then stops (or nearly stops) to feed at a given location for a period of time, and then resumes movement at high speed, stopping again after a distance equal to its length (Figure 5A-E). In this condition, the density profile across the individual’s length at the end of the feeding phase will be very similar to the final density (Figure 5D), so the individual’s feeding rate at the end of the feeding phase nearly matches the marginal feeding rate.

**Figure 5:**
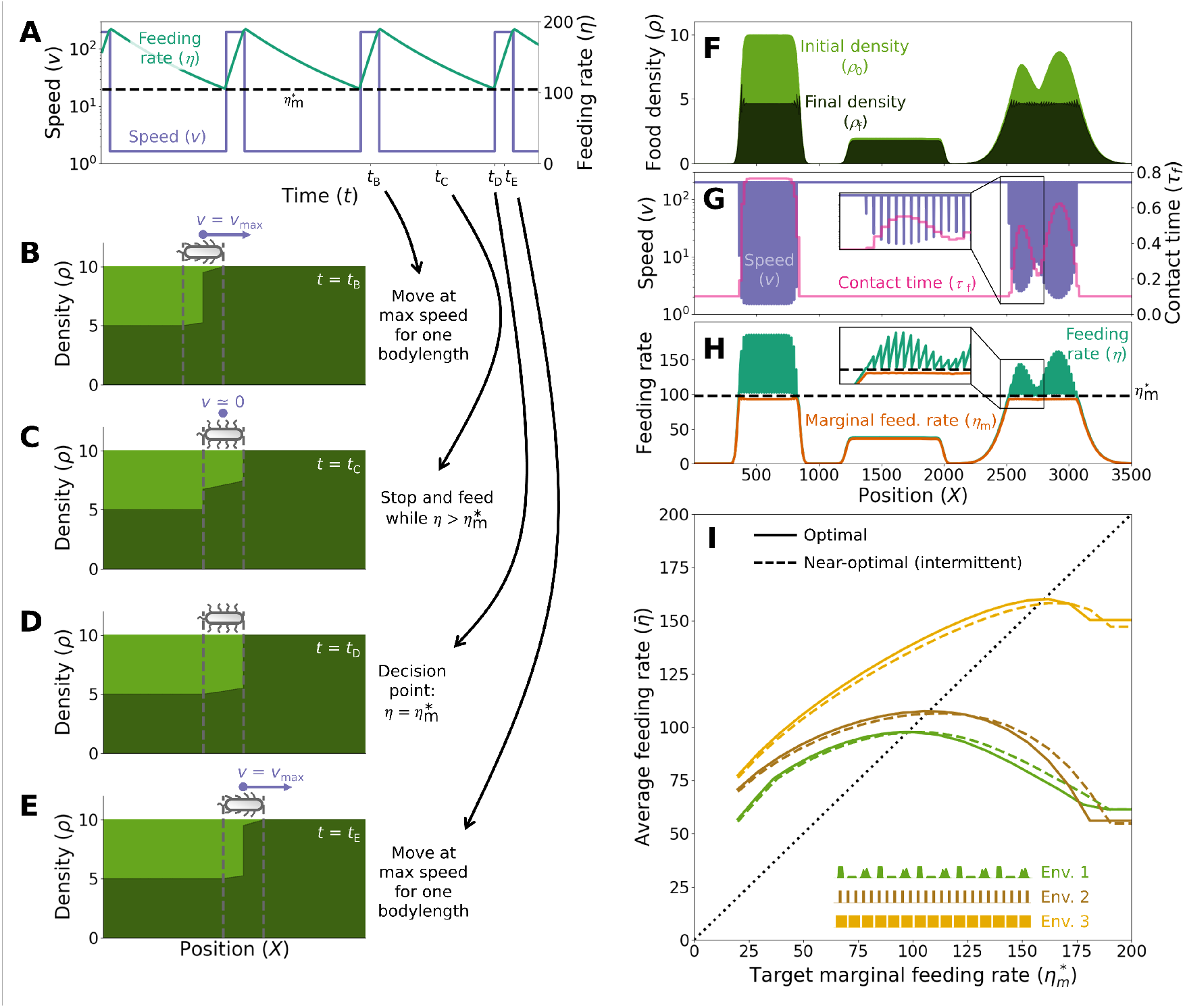
Intermittent movement achieves robust near-optimal behavior. **A**. Speed (*v*, blue) and feeding rate (*η*, green) over time, for an animal performing near-optimal intermittent movement in an environment with constant initial food density. Dashed black line: Target marginal feeding rate 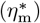. **B**. Food density at each point of space, for the same individual in (A) and at time *t*_*B*_, when the individual is moving at maximum speed. Vertical dashed lines: Region occupied by the individual. **C**. Same as (B), but at time *t*_*C*_, when the individual has stopped to feed. **D**. Same as (B), but at time *t*_*D*_, when the instantaneous feeding rate has just reached the threshold and the individual is about to speed up again. **E**. Same as (B), but at time *t*_*E*_, when the indivdual is moving at maximum speed. **F**. Initial food density (*ρ*_0_, light green) and final food density (*ρ*_f_, dark green) at each point, for an individual implementing the near-optimal intermittent movement. **G**. Speed at every point (*v*, blue) and time that every point spends in contact with the individual (*τ*_f_, pink), for an individual implementing the near-optimal intermittent movement in the environment depicted in (F). Inset: Zoom to the area in the black box. **H**. Feeding rate experienced by the individual at every point (*η*, green), and marginal feeding rate achieved at every point (*η*_m_, orange). Black dashed line: Target marginal feeding rate 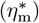. Inset: Zoom to the area in the black box. **I**. Long-term average feeding rate 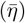 in three different repetitive environments, as a function of the target marginal feeding rate 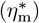. Solid lines correspond to the optimal behavior, and dashed lines to the near-optimal intermittent movement. Inset: Initial density for each of the environments.

Therefore, near-optimal behavior can be achieved by feeding while at low speed until the feeding rate matches the target marginal feeding rate, then moving at maximum speed until the individual does not overlap with its previous feeding spot, and then stopping again to feed (Figure 5A). This behavior achieves a marginal feeding rate equal to the target in the points where the individual stopped and, when food density changes slowly compared to the size of the individual, it also achieves the target feeding rate everywhere else (Figure 5F-H). As a result, it achieves almost the same performance as the optimal behavior (Figure 5I).

Note that implementing this rule requires no knowledge about the depletion dynamics, and no complex computations. The individual only needs to remember one value (the target marginal feeding rate, 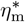 and implement a simple algorithm to achieve optimal performance regardless of the depletion dynamics (Figure S2). While we do not expect any real organism to perfectly implement this intermittent movement, this result shows that variability in speed may help increase foraging efficiency.

## Learning the optimal marginal feeding rate

In order to implement the optimal behavior, the individual must know the optimal target marginal feeding rate 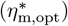, which depends on the environment. As we saw above, this optimal value equals the long-term average feeding rate 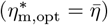. This equality is key, because it means that the optimal behavior can be easily approximated by computing the average feeding rate over a relatively long time in the past. However, this equality is also deceptively simple: The average feeding rate depends on what strategy we are using, and the equality is only true for the optimal strategy. Therefore, an animal that starts with a wrong target marginal feeding rate will get a bad estimate of the optimal target marginal feeding rate.

However, the problem turns out to be easily solvable: For the classical (patch-based) marginal value theorem, even if we start from a value of 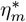 far from the optimum (which will give a behavior far from the optimal one, and therefore an estimate of 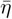 far from the optimal one), adjusting 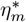 over time with the average of the past feeding rate always converges to the optimal value [28].

We used this result to define a simple learning rule that approximates the average feeding rate and can be easily implemented. This rule corresponds to the differential equation

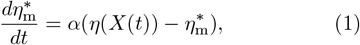

where *α* is a constant that sets the timescale of learning, which must be slow enough to even out the heterogeneity of the environment but fast enough to adapt to new environments. This differential equation leads to the optimal target feeding rate (see Methods for a proof), and is easy to implement even with very simple dynamical systems, such as single neurons or chemical reactions inside a bacterium.

Combining this learning rule with any of our near-optimal control rules leads to near-optimal behavior. Even if the initial estimate of the target marginal feeding rate is incorrect, the estimate eventually converges to the optimal value, and the individual reaches near-optimal performance (Figure 6 and Figure S3).

**Figure 6:**
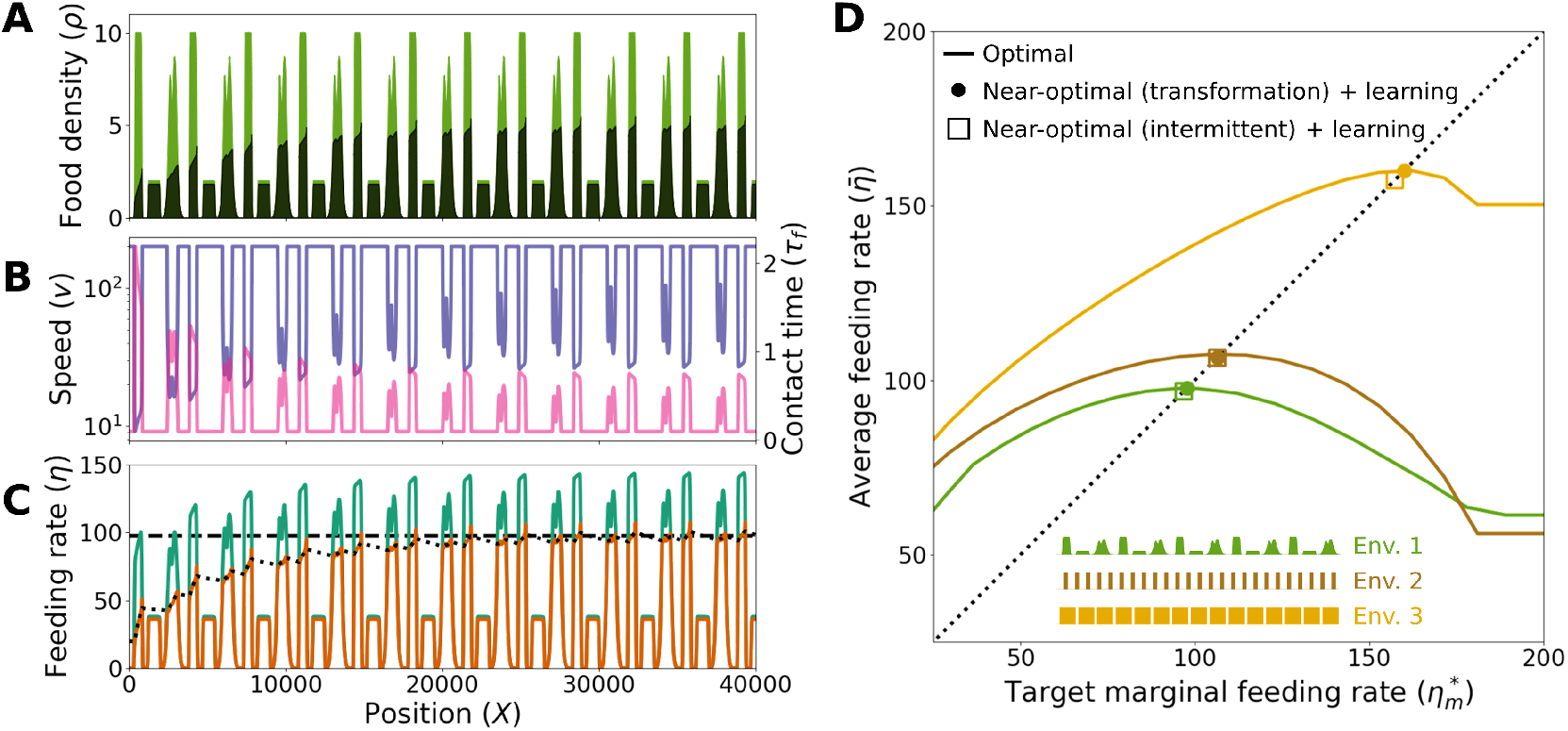
The optimal target marginal feeding rate can be learned. **A**. Initial food density (*ρ*_0_, light green) and final food density (*ρ*_f_, dark green) at each point in a repetitive environment, exploited by an individual implementing learning and the near-optimal behavior explained in Figure 4, which is based on transforming the feeding rate into an estimated marginal feeding rate. **B**. Speed at every point (*v*, blue) and time that every point spends in contact with the individual (*τ*_f_, pink). **C**. Feeding rate experienced by the individual at every point (*η*, green), and marginal feeding rate achieved at every point (*η*_m_, orange). Dotted line: Learned target marginal feeding rate 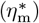, which starts very far from the optimum and evolves according to Equation 1 with *α* = 10^*−*2^. Dashed horizontal line: Optimal value of the target marginal feeding rate 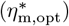. **D**. Long-term average feeding rate 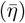 in three different repetitive environments, as a function of the target marginal feeding rate 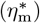. Solid lines: Optimal behavior. Dots: Result of the learning process with the near-optimal behavior explained in Figure 4, which is based on transforming the feeding rate into an estimated marginal feeding rate. Squares: Result of the learning process with the near-optimal behavior explained in Figure 5, which is based on intermittent movement (see Figure S3 for more details).

## Discussion

We have shown that a simple organism with a single sensory input can implement a near-optimal version of the Marginal Value Theorem. This result has four steps: First, we have shown that an animal with no sensory constraints can implement a perfectly optimal behavior by leveling the marginal feeding rate across the environment, and that this strategy works for any arbitrary environment. This generality is important in simple organisms, because they have no way of assessing whether the environment is patchy, as assumed by the classical Marginal Value Theorem. The second step added the constraint of perceiving the environment through a single sensory input (the feeding rate), showing that a near-optimal behavior can still be achieved. However, this near-optimal behavior required knowing the depletion dynamics, so the third step showed that using intermittent movement achieves a more robust near-optimal behavior, which works even when the depletion dynamics are unknown. The fourth step showed that the optimal target marginal feeding rate can be easily learned, since it is equal to the long-term average feeding rate of the environment, as long as the environment is repetitive.

Our model is developed in one dimension because it aims to explain how an organism must change its speed along a given trajectory, and not how this trajectory was chosen. Therefore, one should interpret our space coordinate (*X*) as the distance along a trajectory in a 2D or 3D space, and our initial density as the density the individual finds at each step. If the trajectory intersects itself, the density found at the intersection points will depend on the individual’s previous behavior, but our model can still be applied at every point, and it may still be a good description for a simple organism with limited spatial awareness and memory.

However, a limitation of our work is that a 2D or 3D model may require a more complex analysis to incorporate important realistic features. First, food density in actual environments is usually highly correlated in space, so an organism that encounters a sudden decrease in food density may want to reverse course to return to the high-density region. Our model does not allow the individual to make such decisions, as the trajectory is predetermined. Also, in 2D or 3D the speed may be highly constrained by the turning rate. Finally, re-visits to the same location may dramatically modify the optimal behavior: Our model assumes that every location will be visited only once, so the individual must achieve the target marginal feeding rate the first time it visits every point. However, if a location is going to be re-visited several times, the optimal behavior would require under-exploiting it in the first visits, and reaching the target marginal feeding rate only in the last visit. These considerations are out of the scope of our simple model, but must be taken into account when describing the behavior of real organisms.

We have shown that the optimal strategy will level the marginal feeding rate at every point where the individual is not moving at maximum speed. This is the perfect optimum, before implementing any sensory constraints, and this problem had been studied before, with an almost equivalent conclusion: The optimal behavior should level the final density at every point where the individual was not moving at maximum speed [4]. Our results are equivalent in most situations, and can be considered a re-derivation. However, our formalism has the following advantages:

First, the original formalism required food density to be roughly constant across the length of the individual [4]. While this is a reasonable assumption in most cases, it will necessarily break at the edge of sharp food patches, and [4] did not give an explicit description of the optimal behavior in those situations. In contrast, our formalism defines the optimal behavior at every point, because the marginal feeding rate is well defined everywhere, even when the food density changes abruptly. Note that this generality applies to our general result before adding any sensory constraints; to derive the near-optimal behavior with a single sensory input, we also assumed that density changes slowly in space.

Second, our formalism has an explicit parallel with the classical (patch-based) Marginal Value Theorem (Figure 3). This parallel is made possible by the definition of the marginal feeding rate, which also simplifies the mathematical derivations compared to those used in the original works [2–4].

Third, and most importantly, our formalism shows that a forager can easily learn the relevant statistics of the environment. To implement the optimal behavior, an animal must level the marginal feeding rate at a value equal to the long-term average feeding rate of the environment 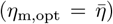. This result is key, because computing this long-term average is very easy in practice (Figure 6). In contrast, the formalism of [4] requires the animal to level the food density down to some critical density, *ρ*_*c*_, but there is no easy way to compute this value, as it is a complex non-linear function of the food density in every location.

A key assumption to derive the equivalence between the optimal target marginal feeding rate and the longterm average feeding rate is that the environment is repetitive. This assumption is not specific to our work, and was already necessary in the original version of the Marginal Value Theorem [13]. In that case, a repetitive environment meant that the expected travel time to the next food patch and the expected quality of the next food patch are the same every time an individual leaves a food patch. To generalize this assumption to arbitrary food distributions, we have assumed that the environment can be divided into many sub-environments, all of which provide the same average feeding rate. This assumption is true, at least approximately, in large environments with stationary food distribution, but it’s important to note that the same result can be derived even if the environment does not fulfill this condition, from an argument about uncertainty: If we assume that the individual has no information about the density it will encounter next, and that the environment has very little spatial correlation, then the expected value of the feeding rate for the next small portion of the environment is equal to the long-term feeding rate. This fact, together with our Lemma 1 (see Methods), means that the individual should move to this new portion of the environment leaving behind a marginal feeding rate equal to the average feeding rate.

Another limitation of our study is that the pattern of intermittent movement that implements robust near-optimal behavior requires the animal to move by exactly one body length at a time and then abruptly slow down (Figure 5). This ideal pattern is more stereotyped than movement patterns found in most real organisms, but other movement patterns may also achieve a similar outcome, and it’s important to note that any variability in speed may be helpful to get a more robust estimate of the marginal feeding rate.

While our model should be applicable to a wide range of simple organisms, it may be useful to discuss how it applies to a particular one, which has inspired much of our work: The nematode *Caenorhabditis elegans. C. elegans* is a small worm (1 mm long), with a simple nervous system (around 300 neurons in total) [15, 21]. It has no vision or audition, so it has very limited spatial awareness. Despite these sensory limitations, *C. elegans* successfully chooses the densest food patches when given several choices [26, 39]. *C. elegans* modulates its speed in response to food, slowing down when it finds high-density food [6, 16, 23, 38], and leaves food patches once they are depleted [30]. Also, it remembers the food density experienced in the past, and modulates its response to food accordingly, in a way that is consistent with our learning rule, at least qualitatively [1, 38]. It feeds on bacteria, which at intermediate and high concentrations can be described as a continuous food source, rather than as discrete prey items. It can sense bacteria outside its body through mechanical and chemical sensing [5, 19] and it is likely that these stimuli are important to react to changes in food density, so it probably uses more than one sensory input. However, it is reasonable to think that feeding rate is its main way of estimating food density: It feeds by pumping bacteria into its pharynx, and its pumping rate is proportional to food density, even in laboratory conditions where the worm is restrained inside a capillar with its external sensory stimuli severely affected [25]. For these reasons, *C. elegans* provides an example where the assumptions of our model are at least approximately met. We expect many other simple organisms to also meet our assumptions.

In conclusion, our study extends the Marginal Value Theorem to organisms with minimal sensory and cognitive capacities, broadening its applicability beyond the animal species typically studied in Ethology. By bridging theoretical optimality with practical constraints, our findings unify our understanding of foraging strategies across the tree of life. This approach complements existing studies on more complex foragers and sets the stage for future exploration of adaptive behaviors in simple organisms, emphasizing their evolutionary ingenuity and ecological significance.

### Proof that the optimal behavior levels the marginal feeding rate

#### Main Theorem

Let *v*(*X*) be the optimal strategy in a given environment. Then, the optimal strategy must satisfy that

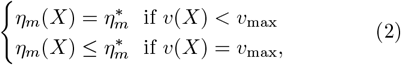

where 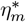 is a constant that we call *target marginal feeding rate*.

**Proof:** We will prove this result by contradiction.

First, note that Equation 2 is equivalent to saying that for any point *X*_1_ with *v*(*X*_1_) *< v*_max_, and for any point *X*_2_, we must have *η*_*m*_(*X*_1_) ≥ *η*_*m*_(*X*_2_).

Now consider an optimal strategy *v*(*X*) that breaks this statement, so it has had at least two points, *X*_1_ and *X*_2_, with *v*(*X*_1_) *< v*_max_, and with *η*_*m*_(*X*_1_) *< η*_*m*_(*X*_2_).

Then, we can define an alternative strategy, *v*^*′*^(*X*), which is identical to *v*(*X*) except that it has an infinitesimal increase of speed to spend Δ*t* less time at *X*_1_ and an infinitesimal decrease of speed to spend Δ*t* more time at *X*_2_. These infinitesimal changes in speed result in the individual eating Δ*t η*_m_(*X*_1_) less food around *X*_1_ and Δ*t η*_m_(*X*_2_) more food around *X*_2_, where *η*_m_ is the marginal feeding rate (see the proof in Lemma 1). Therefore, the total amount of food collected by strategy *v*^*′*^(*X*) is

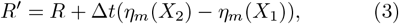

where *R* is the amount of food collected by the optimal strategy. Given that *η*_*m*_(*X*_1_) *< η*_*m*_(*X*_2_), we have that *R*^*′*^ *> R*, which is impossible, because *R* is the amount of food collected by the optimal strategy. Q.E.D.

##### Lemma 1

Decreasing the speed at point *X* during an infinitesimal amount of time Δ*t* increases the total amount of food *R* by Δ*t η*_m_(*X*). Likewise, an equivalent increase of speed decreases *R* by the same amount.

**Proof:** The first step of the proof is to find a more explicit functional form for the food density and depletion rate at a given point *x*. When a point *x* is in contact with the individual, its density *ρ* is governed by the differential equation

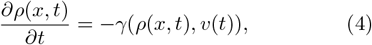

where *γ* is the depletion rate, which, as discussed in the main text, depends on the food density and on the individual’s speed. The solution of this differential equation may be very complicated, especially if the speed changes over time. However, for a given point *x* we only need to consider the relatively brief time interval in which the point is in contact with the individual. We make the simplifying assumption that speed changes sufficiently slowly so that we can neglect the effect of speed changes in Equation 4. We can then write

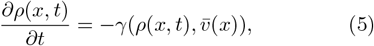

where 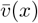 is the average speed of the individual during the time it was in contact with point *x*. Now we note that Equation 5 is only valid while point *x* is in contact with the individual, with food density at *x* remaining constant whenever the individual is not in contact with it. It is therefore convenient to perform the variable change *τ* = *t − t*_*x*_, where *t*_*x*_ is the time when the individual touches point *x* for the first time (i.e. the time when the individual reaches *x − r*). Then we can write

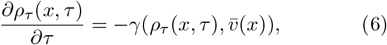

where we have added sub-index *τ* to distinguish the mathematical function with explicit dependence on *τ* from the one with explicit dependence on absolute time *t*. Equation 6 is valid for *τ* ∈ [0, *τ*_f_(*x*)], where *τ*_f_(*x*) is the total time that point *x* spends in contact with the individual. Because our proof must remain valid for any shape of the depletion rate *γ*(*ρ, v*), we cannot give a closed solution for Equation 6, and in some cases no analytical solution will exist. However, given that 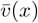 is constant over time, we can ensure that the solution to this equation will have the form

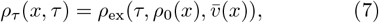

where *ρ*_0_(*x*) is the initial density at *x*. We have added the subindex “ex” to distinguish *ρ*_ex_ as a mathematical function that depends explicitly on *τ, ρ*_0_ and 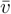. Given that *ρ*_0_(*x*) and 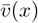 are constant in time, this equation means that the food density at *x* only depends on the time that the individual has spent in contact with *x*, regardless of its previous history. This property will become important later on.

We can now compute the total amount of food eaten by the individual as

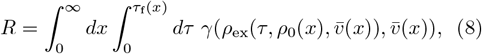

where *τ*_f_(*x*) is the total time the individual spends in contact with point *x* (note that *τ*_f_(*x*) = 0 for points that the animal does not reach in the allotted time, so the spatial integral is zero outside the region traversed by the individual).

Let us now consider an alternative strategy, *v*^*′*^(*X*), which produces the alternative schedule 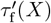. This strategy differs from the original one only in an infinitesimal speed bump: At *X*_1_ the individual speeds up during an infinitesimal distance, and then goes back to the original strategy. This speed bump translates in the individual spending Δ*t* time less at *X*_1_, where Δ*t* is an infinitesimal amount of time. Therefore, every point between *X*_1_ and *X*_1_ + *r* spends Δ*t* time less in contact with the individual (see Lemma 2 for a formal proof), and the new strategy is

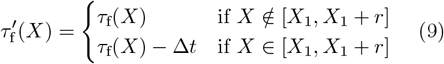

Therefore, the amount of food eaten with the alternative strategy is

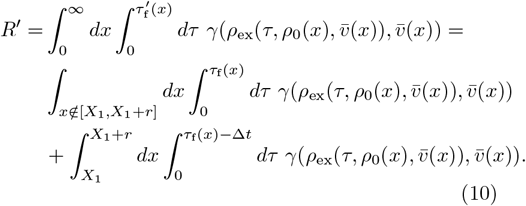

We can now add and subtract the term 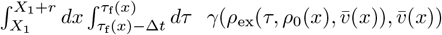 getting

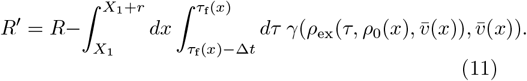

Now, since Δ*t* is infinitesimal, we can solve the time integral as if 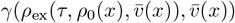 was constant during Δ*t*, getting

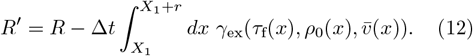

Now we define explicitly the marginal feeding rate (*η*_m_), as the feeding rate the individual would experience if it came back to an already-visited point, so

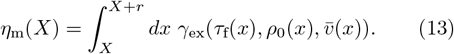

And therefore,

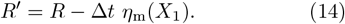

This shows the lemma for an infinitesimal increase in speed, and the same process can be followed to show that a strategy *v*^*′′*^(*X*) that has an infinitesimal decrease in speed at *X*_2_ will collect the amount of food *R*^*′′*^ = *R* + Δ*t η*_m_(*X*_2_).

##### Lemma 2

Given a strategy *v*(*X*) and its associated schedule *τ*_f_(*X*), increasing the speed at point *X*_1_ during an infinitesimal distance results in a new schedule 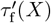 such that

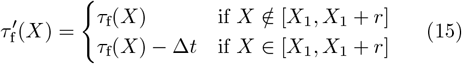

**Proof:** Consider a strategy *v*(*X*) and its associated schedule *τ*_f_(*X*). Then consider the alternative schedule

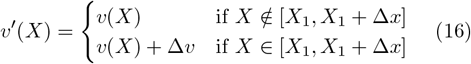

where Δ*x* is infinitesimal. Now we distinguish three types of points (see Figure S4):

First, points unaffected by the speed bump, which are those where *X* ∉ [*X*_1_, *X*_1_ + *r* + Δ*x*] (green in Figure S4). For these points, we simply have 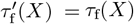.

Second, points fully affected by the speed bump, which are those where *X* ∈ [*X*_1_+Δ*x, X*_1_+*r*] (red in Figure S4). To compute the new schedule in these points, we must compute the amount of time that each point is in contact with the individual. At constant speed this time would be *r/v*. In general, this time is

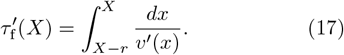

For a point affected by the speed bump (i.e. *X* ∈ [*X*_1_, *X*_1_ + *r*]), we can split the integral in Equation 17 as

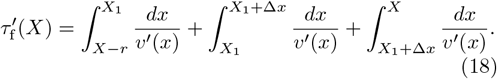

Using the definition of the new strategy for each period, we get

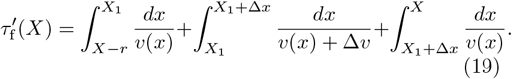

Now we add and subtract the term 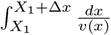, getting

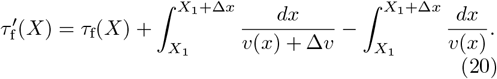

Given that these integrals are over an infinitesimal segment Δ*x*, we can solve them as if *v*(*x*) remains constant over them, so we get

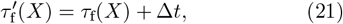

where

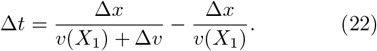

Finally, we must consider the points that are partially affected by the speed bump. These are points for which *X* ∈ [*X*_1_, *X*_1_ + Δ*x*] or *X* ∈ [*X*_1_ + *r, X*_1_ + *r* + Δ*x*] (orange in Figure S4). However, these points only cover two infinitesimal intervals of size Δ*x*. Given that all our calculations involve integrals covering at least the individual’s length *r*, which is much bigger than Δ*x*, we can neglect these points and we can approximate the new schedule by Equation 15. Q.E.D.

### Proof that the optimal target marginal feeding rate equals the average feeding rate

Now we will prove that the optimal value for 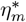 is equal to the long-term average feeding rate 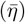. To do this, we need to add an extra assumption, called “repetitive environment”. In the original MVT paper, this assumption stated that the expected travel time to a new patch was constant over time [13]. The equivalent assumption in our case is that the environment is long and repetitive, so it can divided in many (ideally infinite) sub-environments that provide the same average feeding rate. Let us now consider an environment with such properties, and its associated optimal strategy that levels the marginal feeding rate at 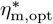 and achieves an average feeding rate 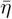. We can subdivide this environment into many sub-environments, all with the same average feeding rate 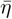.

We will now consider an alternative strategy that will go infinitesimally faster, saving enough time to lengthen the trajectory and cover a new sub-environment. Let *t*_sub_ be the time needed to cover this new sub-environment achieving the average feeding rate 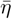 (and therefore extracting an amount of food 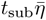 from this sub-environment). The alternative strategy has an infinitesimally higher target marginal feeding rate, 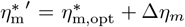, which makes the individual go infinitesimally faster in every point where its speed is lower than the maximum, saving a total time *t*_sub_. Note that given that the trajectory is very long, even if the change in speed at any given point is infinitesimal, *t*_sub_ adds up to a finite amount of time. But given that the change in speed at any point is infinitesimal, we can use Lemma 1 to compute the amount of food that the individual does not eat because of the speedup as 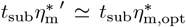. Therefore, the amount of food collected by this new strategy is

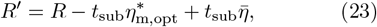

where *R* is the amount of food collected by the optimal strategy.

Similarly, we can consider another equivalent strategy where the animal slows down infinitesimally, shortening the trajectory by one sub-environment. In this case, the amount of food collected is

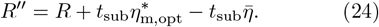

Given that *R* was the amount of food collected by the optimal strategy, we must have *R*^*′*^ ≤ *R* and *R*^*′′*^ ≤ *R*. Therefore, from Equations 23 and 24, we must have that

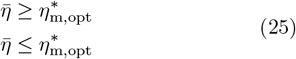

As both of those conditions must be true at the same time, it follows that 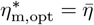. Q.E.D.

### Near-optimal behavior with a single sensory input

Here we show how an individual whose only sensory input is the feeding rate *η* can estimate its marginal feeding rate *η*_m_. To perform this estimation, we assume that the initial food density (*ρ*_0_) and the speed (*v*) change slowly at the scale of the individual’s length. Therefore, we can assume that *ρ*_0_ and *v* are constants. In this situation, the individual always depletes the food to the same final density (Figure 4A). This final density depends on the initial density and on the individual speed, so we call it *ρ*_f_(*ρ*_0_, *v*). This function depends on the shape of the depletion function *γ*(*ρ, v*), and the results of this section depend heavily on the exact shape of this function. We will first develop the particular case of exponential depletion, which is the one we use to illustrate our results in the figures. Then, we will show the more general derivation.

#### A particular case: Exponential depletion

Here we assume that depletion rate does not depend on speed and is simply proportional to density,

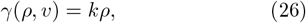

where *k* is a constant. Therefore, density at every point in contact with the individual is governed by the differential equation 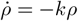, whose solution is

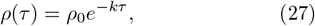

where *τ* is the time that the point spent in contact with the individual. Therefore, in this particular case the density at every point decreases exponentially with the time spent in contact with the individual.

Given that the individual spends a total time of *r/v* in contact with any given point, the final density in this particular case is

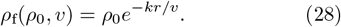

Since the final density is roughly constant in space across the section, the integral for the marginal feeding rate (Equation 13 can be solved easily, giving

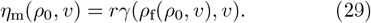

Substituting here the expressions for the depletion rate (Equation 26) and the final density (Equation 28), we get

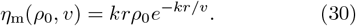

The individual knows its speed *v* but it does not know the initial density, *ρ*_0_. Therefore, the next step is to estimate this initial density from the information available to the individual. To do this, we first note that when *ρ*_0_ and *v* are constant, the feeding rate can be simply calculated as the amount of food that was initially available across the individual’s length (*rρ*_0_), minus the amount of food left after the individual passes (*rρ*_f_(*ρ*_0_, *v*)), over the time it takes to pass (*rv*). Therefore,

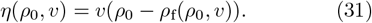

Substituting here the expression for the final density (Equation 28), we get

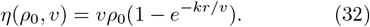

We can solve for *ρ*_0_ in this equation, getting *ρ*_0_ as a function of *η* and *v*,

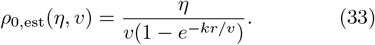

We call this function *ρ*_0,est_(*η, v*), because it represents the initial density that the individual estimates from its feeding rate and its speed. Using this estimate and the target marginal feeding rate, we can solve Equation 30 to get the target speed,

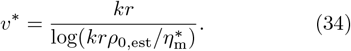

To perform the transformations shown in Figure 4, we can substitute *ρ*_0_ by the estimated initial density into the marginal feeding rate (Equation 30), to get an estimated marginal feeding rate,

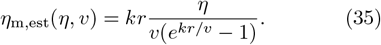

Finally, we can solve for *η* in order to get the target feeding rate that the animal must achieve at each speed (*η*\*(*v*), red line in Figure 4C):

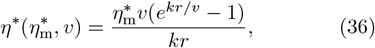

where 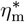 is the target marginal feeding rate.

#### General case

The general case follows the same steps as the particular case above. We assume that the functions *γ*(*ρ, v*) and *ρ*_f_(*ρ*_0_, *v*) are known, but we do not specify their exact formula. The general expression of the marginal feeding rate in Equation 29 is still valid, as is the expression for the feeding rate in Equation 31. Assuming that we can solve for *ρ*_0_ in this equation, we can compute the estimated initial density, *ρ*_0,est_(*η, v*). Then, the estimated marginal feeding rate is given by

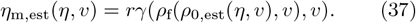

Finally, we must solve for *η* in this equation to obtain the target feeding rate at any speed for a given value of the target marginal feeding rate,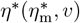.

### Alternative depletion dynamics

For most of the simulations in this manuscript, we assume simple exponential depletion dynamics, described by Equations 26 and 27. However, in Figures S1 and S2, we assume different depletion dynamics to test the robustness of each strategy. The depletion dynamics used in these two figures assume that depletion rate is proportional for low densities but saturates when reaching some threshold density (*ρ*_s_),

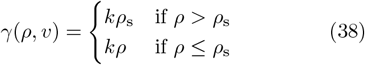

The density at every point in contact with the individual is governed by the differential equation 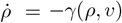. The solution of this differential equation is

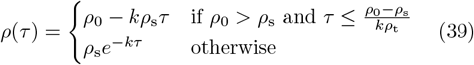

### Learning converges to the optimal target marginal feeding rate

We start from the assumption of a repetitive environment, which states that the environment can be subdivided into segments, each giving the same average feeding rate. We then use averaging theory [37, 43], which says that a differential equation of the form

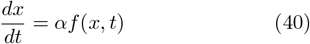

where *f* (*x, t*) is periodic in time and *α* is sufficiently small, can be approximated by

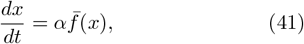

where

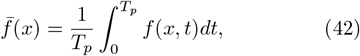

where *T*_*p*_ is the duration of one period. In our case, *T*_*p*_ is the time it takes the individual to cross one of the repetitive segments of the environment. Applying this result to Equation 1, we get

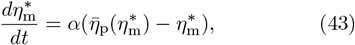

where 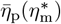 is the average feeding rate experienced in one segment of the environment, when using the threshold 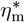.

We know that the optimal value of 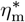 fulfills 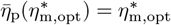 (see for example Figure 2D). Therefore, Equation 43 has a fixed point at 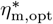. To show that this fixed point is stable, we linearize the system around the fixed point and require that a perturbation decay with time (i.e., that it has a negative eigenvalue). This leads to the condition

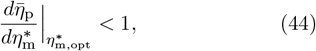

which is automatically satisfied since in our case

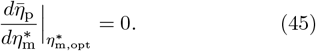

### Discrete model

#### Description of the model

To implement the model and produce simulations, we built a discrete version of it. The space is divided into discrete bins, which contain a certain density of food. The section representing the individual covers 20 bins in all simulations, and the position of the animal is considered to be the first bin of the section.

The model iterates in space: The individual has a defined speed in each iteration, whose inverse is the time it spends before advancing by one bin. During this time, the individual depletes food from all bins in contact with it, and we compute analytically the amount of food eaten from each bin using Equation 27. The feeding rate of the individual (*η*) is the sum of the food eaten across all the bins in contact with the individual, divided over the time spent in the current iteration. We then move to the next iteration, and the simulation stops when the sum of the times in all iterations (i.e the total time) reaches *T*, the limit time of the simulation. To compute the marginal feeding rate in every point, we first run the full simulation to find the final density in every bin. Then, for every position in space we compute the depletion rate of every bin in contact with the individual using Equation 26, and we sum across all the bins in contact with the individual.

#### Finding the optimal strategy in the discrete model

To find the strategy that perfectly levels the marginal feeding rate in a given environment, we proceeded as follows. In our discrete model, the strategy is defined by the amount of time the individual spent in every bin (where the individual’s position is defined at the last bin it covers), which is equal to the inverse of the individual’s speed. We started from a strategy that has the maximum speed at every point (i.e. it spends the minimum possible amount of time at every point). Then, we computed the final density at every bin after running this strategy, and the marginal feeding rate at every point resulting from that final density. We ignored the points where the individual moved at maximum speed and the marginal feeding rate was lower than the target marginal feeding rate, because these points already fulfill the conditions for the optimal behavior (see Equation 2). For the remaining points, we computed the difference between the marginal feeding and the target marginal feeding rate, and we selected the point where the absolute value of this difference was maximal. Then, we increased or reduced the amount of time the individual spent in that point by a time increment (which initially was equal to 0.05), in order to bring the marginal feeding rate closer to the target marginal feeding rate. We then ran the new strategy, computed the final density at every bin, and the associated marginal feeding rate at every point. We computed the difference between the target marginal feeding rate and the closest marginal feeding rate from it among bins in which it is possible to relocate time. If this value is lower than at the previous step, we accepted the new strategy and repeated the whole procedure. Otherwise, we rejected the new strategy, reduced the value of the time increment by 1%, and repeated the whole procedure. We iterated this process until the highest absolute difference between the marginal feeding rate and the target feeding rate was lower than 1, for all points where the individual is not moving at maximum speed.

This algorithm produces a strategy that levels the marginal feeding rate in every point where the individual is not moving at maximum speed, fulfilling the conditions for optimality. However, there may exist many different strategies that achieve this result. All of them achieve the same performance (i.e. the same average feeding rate across the whole environment), but they may look very different. The algorithm described in the previous paragraph tends to produce strategies with quick changes in speed, which are not the most realistic. To alleviate this issue we proceeded as follows. Once an optimal strategy was found, we smoothed it by applying a moving average to the time in every point, with a window size of 20 bins. We thus obtained a smoother strategy, but which was not perfectly optimal any more. We then applied again the procedure described in the previous paragraph, starting from this smoothed strategy, to turn it into a perfectly optimal one. We repeated 30 rounds of smoothing plus optimization, which gave us a reasonably smooth optimal strategy, such as the one shown in Figure 2B.

#### Single-input near-optimal behavior in the discrete model

We implemented the near-optimal strategy shown in Figure 4 as follows. The individual starts with an initial speed, which determines the time for the first iteration. Then, each iteration we compute the individual’s feeding rate using the procedure described in Section “Description of the model”. Then, we used this feeding rate and Equation 33 to get an estimate of the initial density (*ρ*_0,est_), and we used this estimate in Equation 34 to get the optimal speed. We then use this speed to determine the time of the next iteration, and repeat the procedure. This method introduces a delay in the computation of speed since it is using values from the previous iteration, but the impact of such a delay is negligible in our conditions.

#### Intermittent movement in the discrete model

The intermittent movement shown in Figures 5 and 6 uses the following algorithm: The individual moves at maximum speed across a distance equal to its length (in our case, 20 bins). Then, it stops to feed, remaining in its current position until its instantaneous feeding rate falls below 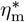 (if the individual’s feeding rate is lower than 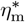 from the beginning, then it will spend as little time as possible, meaning that it will keep moving at maximum speed). After this time the individual moves again at maximum speed across a distance equal to its length, and repeats the process.

#### Learning 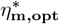 in the discrete model

To produce the results shown in Figure 6, we ran the discrete model with intermittent movement as described in the previous section, but we changed the value of 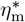 in every iteration, as follows: We started from an initial value of 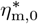, equal to 20 (we chose this initial value because it’s far from the optimum). Then, in every iteration we updated 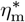 using the formula

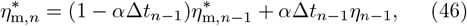

where *α* is a constant that controls the learning rate, 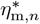 is the value of 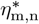 used in the *n*-th iteration, and *η*_*n*_ is the feeding rate experienced by the individual in the *n*-th iteration. Δ*t*_*n*_ is the time spent in the *n*-th iteration, so Δ*t*_*n*_ = 1*/v*_*n*_. We must multiply the learning rate *α* times Δ*t* because our discrete model iterates in space rather than in time, but the average feeding rate must be computed as an average in time. Therefore, the weight of each iteration on the average must be proportional to the time spent in that iteration.

To produce the results shown in Figure S3, we used the same procedure but using the model described in the section “Single-input near-optimal behavior in the discrete model”.

## Code availability

The code used to generate all results and figures of this paper can be found at https://github.com/tomO-L/Optimal-foraging-for-simple-organisms.

## Glossary

*t*: Time.
*T*: Total time.
*x*: Space coordinate.
*X*: Space coordinate.
*r*: Length of the individual.
*τ* (*X, t*): Time that point *X* has spent in contact with the individual by time *t*.
*τ*_f_(*X*): Total time that point *X* spends in contact with the individual.
*v*(*X*): Speed of the individual at point *X*.
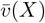: Average speed of the individual while it was in contact with point *X* (i.e. when its position was between *X− r* and *X*).
*ρ*(*X, t*): Food density at point *X* and at time *t*. Units: Amount of food per unit of distance.
*ρ*_0_(*X*): Initial food density at point *X. ρ*_0_(*X*) = *ρ*(*X*, 0). Units: Amount of food per unit of distance.
*ρ*_f_(*X*): Final food density at point *X. ρ*_f_(*X*) = *ρ*(*X, T*). Units: Amount of food per unit of distance.
*ρ*_*τ*_ (*X, τ*): Food density at point *X*, when it has spent a time *τ* in contact with the individual. Units: Amount of food per unit of distance.
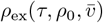: Food density for a point whose initial density was *ρ*_0_, and that has spent a time *τ* in contact with the individual, whose average speed during this time was 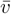. Units: Amount of food per unit of distance.
*γ*(*ρ, v*): Depletion rate at a point whose density is *ρ* and which is in contact with the individual moving at speed *v*. Definition: *γ* = − *∂ρ/∂t*. Units: Amount of food per unit of distance and per unit of time.
*η*(*X*): Feeding rate of the individual when it’s at point *X*. Definition: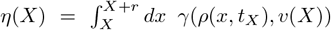, where *t*_*X*_ is the time at which the individual is at *X*. Units: Amount of food per unit of time.
*η*_m_(*X*): Marginal feeding rate at point *X*. Definition: 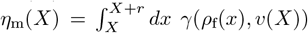. Units: Amount of food per unit of time.
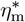: Target marginal feeding rate. Units: Amount of food per unit of time.
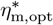: Optimal target marginal feeding rate. Units: Amount of food per unit of time.
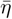: Average feeding rate over the total time *T*. Units: Amount of food per unit of time.
*R*: Total amount of food eaten. Units: Amount of food.

## Acknowledgments

Color schemes for the figures were chosen with https://colorbrewer2.org/ This work was funded by grant ANR-22-CE02-0002 (ForAnInstant) from the Agence Nationale de la Recherche (ANR).

## Author contributions

TOL: Methodology, Software, Formal analysis, Investigation, Writing - Original Draft. RLC: Methodology, Validation, Investigation. JF: Conceptualization, Methodology, Supervision, Writing - Review & Editing. APE: Conceptualization, Methodology, Validation, Formal analysis, Writing - Original Draft, Writing - Review & Editing, Visualization, Supervision, Project administration, Funding acquisition

## Supplementary Information

### Supplementary Figures

**Figure S1:**
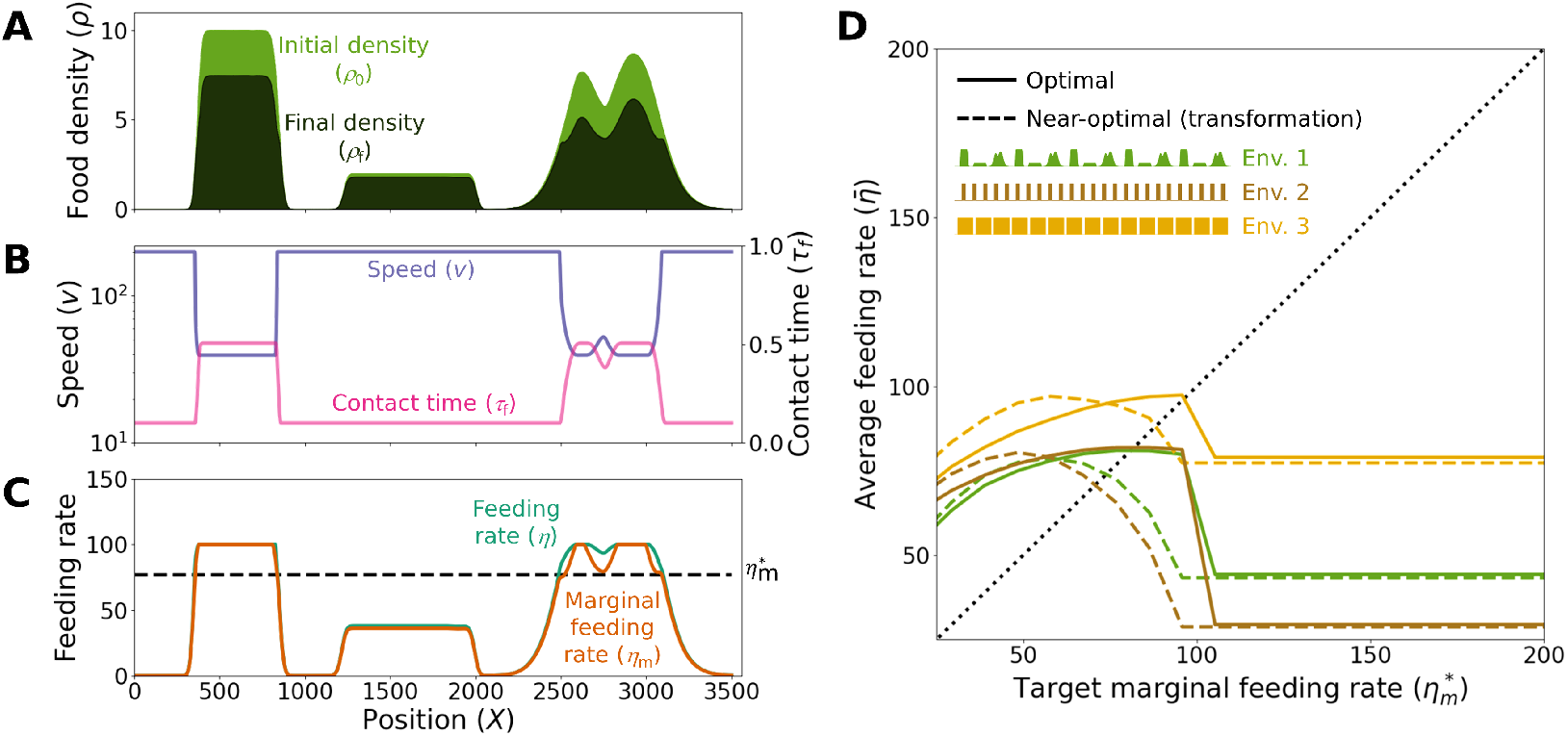
The near-optimal behavior based on estimating the marginal feeding rate from the feeding rate breaks down when depletion dynamics are unknown. In this simulation, the individual is performing the near-optimal behavior described in Figure 4 of the main text, which is based on transforming the feeding rate (*η*) to estimate the marginal feeding rate 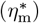. This transformation is made using Equations 35 and 36, which assume that depletion dynamics are exponential (Equations 26 and 27). However, the actual depletion dynamics have saturation at high densities, and are therefore governed by Equations 38 and 39. Because of this mismatch, the near-optimal behavior does not work in this environment. **A**. Initial food density (light green) and final food density (dark green) at each point along a trajectory in a heterogeneous environment. **B**. Speed (blue) and time in contact (purple) for the situation described in A. **C**. Feeding rate (blue) and marginal feeding rate (purple) achieved in every point for the situation described in A. Note that the marginal feeding rate is not leveled at the correct value, and is not even constant in the region of the space where density changes smoothly. **D**. Long-term average feeding rate 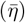 in three different repetitive environments, as a function of the target marginal feeding rate 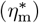. Solid lines: Optimal behavior with no sensory constrains and using the correct shape for the depletion dynamics. Dashed lines: Near-optimal behavior using the wrong shape of the depletion dynamics (as described for panel A). Inset: A portion of the environment represented by each color.

**Figure S2:**
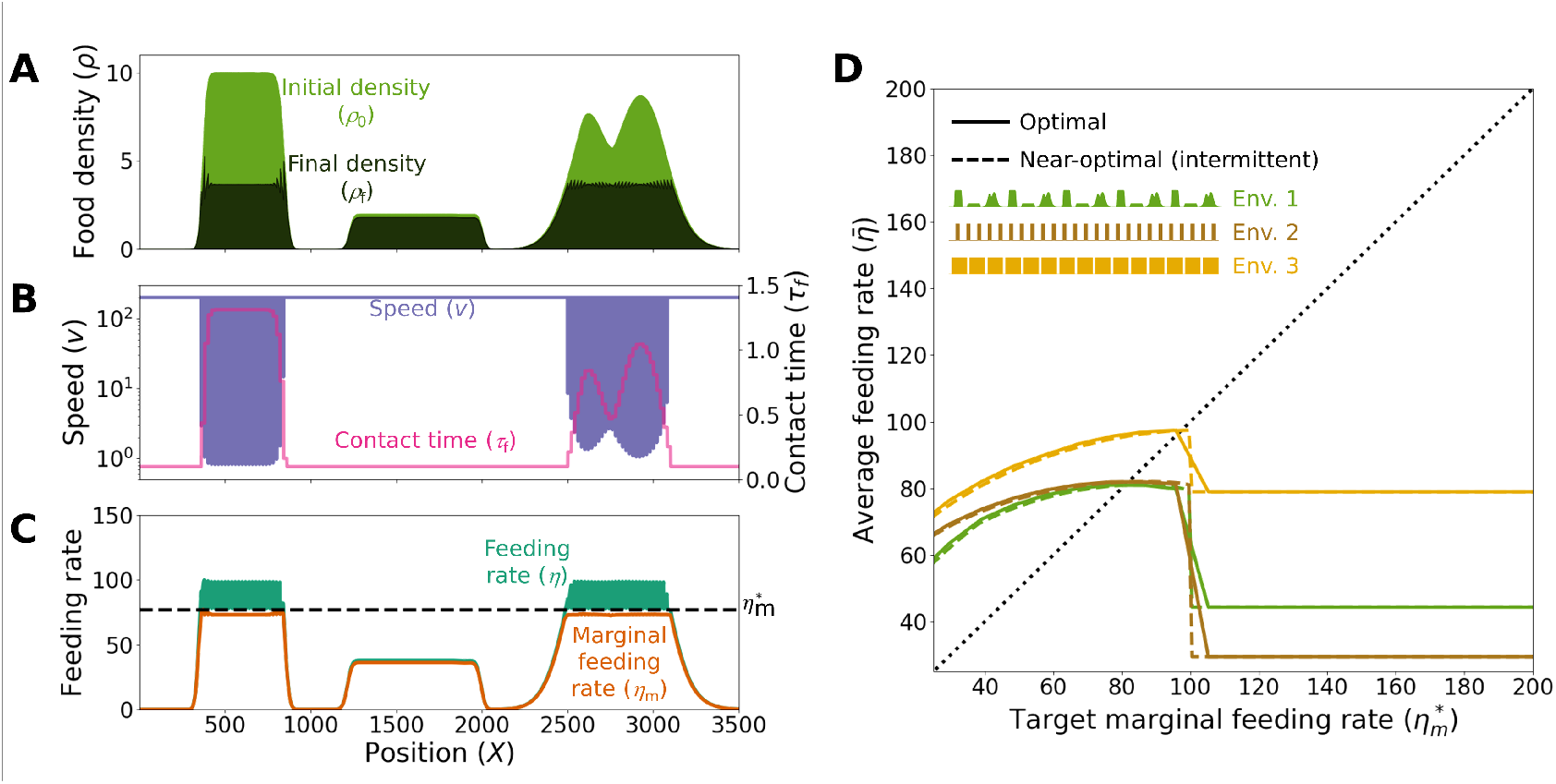
The near-optimal behavior based on intermittent movement works even when depletion dynamics are unknown. In this simulation, the individual is performing the near-optimal behavior described in Figure 5 of the main text, which is based on intermittent movement. The depletion dynamics in this figure have saturation at high densities, and are therefore governed by Equations 38 and 39, but the behavioral rule is independent of the depletion dynamics (and is therefore identical to the one used in Figure 5 of the main text). **A**. Initial food density (light green) and final food density (dark green) at each point along a trajectory in a heterogeneous environment. **B**. Speed (blue) and time in contact (purple). **C**. Feeding rate (blue) and marginal feeding rate (purple) achieved in every point. **D**. Long-term average feeding rate 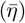 in three different repetitive environments, as a function of the target marginal feeding rate 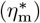. Solid lines: Optimal behavior with no sensory constrains and using the correct shape for the depletion dynamics. Dashed lines: Near-optimal behavior. Inset: A portion of the environment represented by each color.

**Figure S3:**
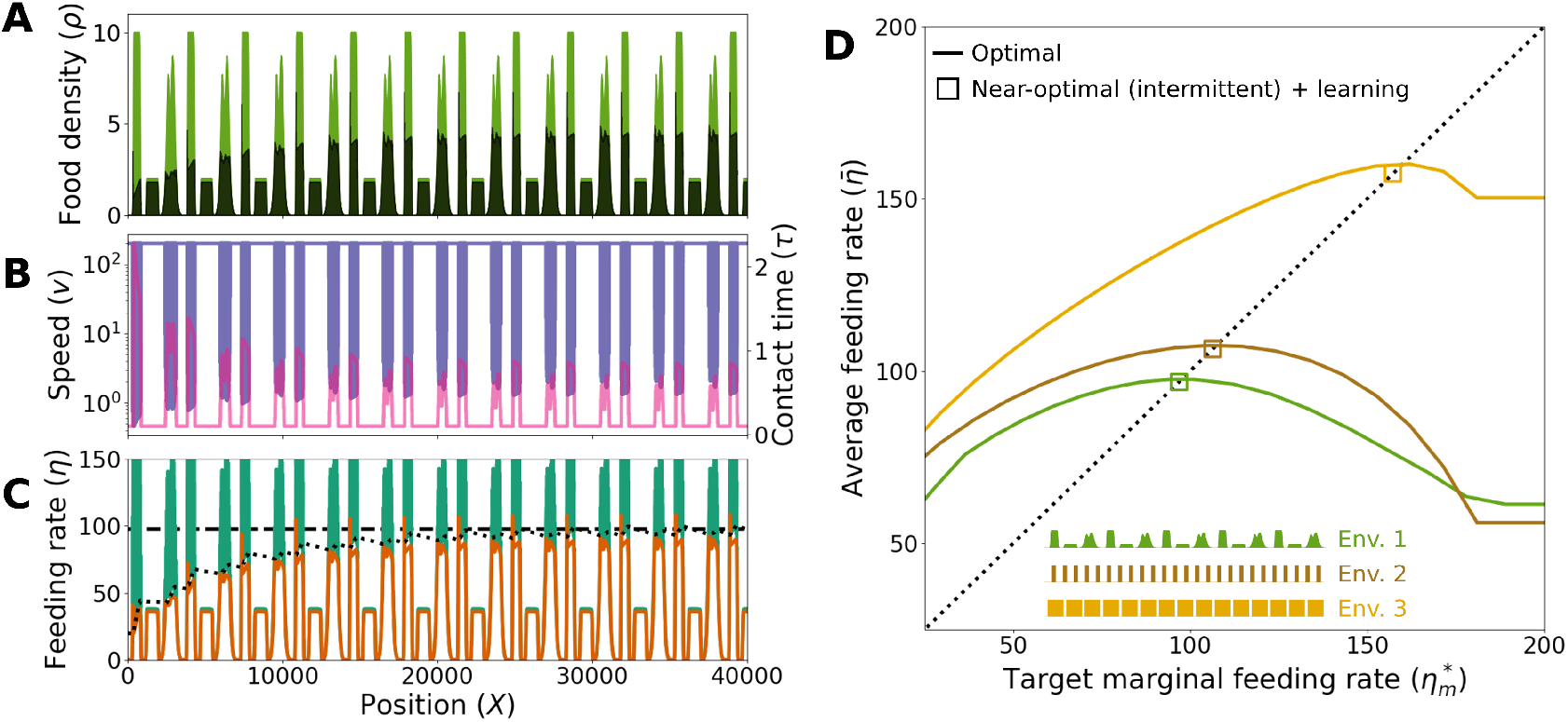
Learning with intermittent movement. **A**. Initial food density (*ρ*_0_, light green) and final food density (*ρ*_f_, dark green) at each point in a repetitive environment, exploited by an individual implementing learning and the near-optimal behavior explained in Figure 5, which is based on intermittent movement. **B**. Speed at every point (*v*, blue) and time that every point spends in contact with the individual (*τ*_f_, pink). **C**. Feeding rate experienced by the individual at every point (*η*, green), and marginal feeding rate achieved at every point (*η*_m_, orange). Dotted line: Learned target marginal feeding rate 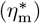, which starts very far from the optimum and evolves according to Equation 1 with *α* = 10^*−*2^. Dashed horizontal line: Optimal value of the target marginal feeding rate 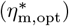. **D**. Long-term average feeding rate 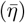 in three different repetitive environments, as a function of the target marginal feeding rate 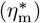. Solid lines: Optimal behavior. Squares: Result of the learning process.

**Figure S4:**
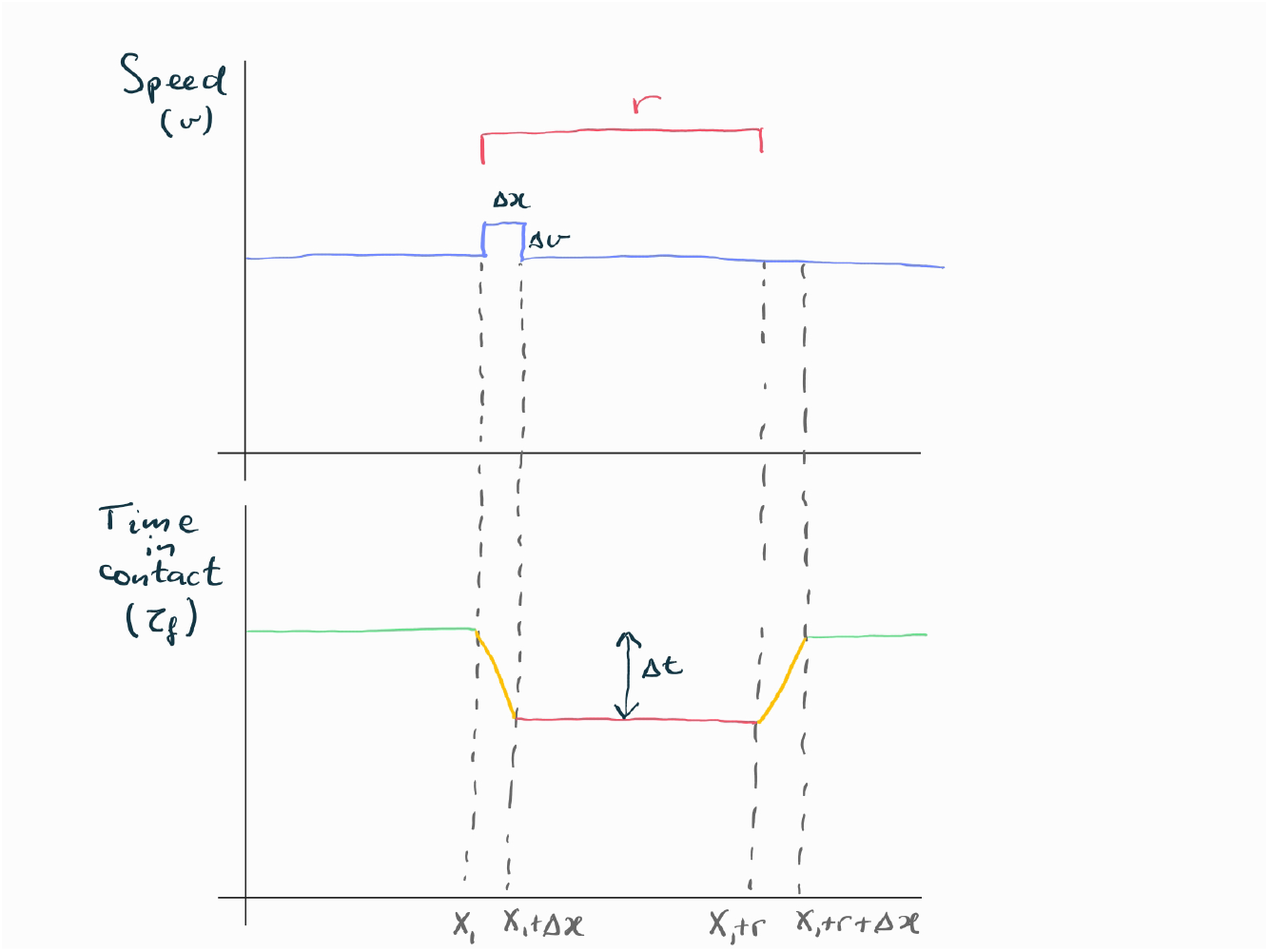
Effect of a speed bump in the time that each point spends in contact with the individual. **A**. Individual’s speed at each point. Speed is constant except for a speed bump of size Δ*v* that happens at *X*_1_ and lasts for an infinitesimal length Δ*x*. The individual has size r (red segment). **B**. Time that each point spends in contact with the individual. Green: Regions not affected by the speed bump. Red: Region fully affected by the speed bump. Orange: Regions partially affected by the speed bump. The orange regions are infinitesimally small and they can be neglected, so the effect of the speed bump is can be approximated by a square decrease of Δ*t* between *X*_1_ and *X*_1_ + *r*.

